# Bias in two-sample Mendelian randomization when using heritable covariable-adjusted summary associations

**DOI:** 10.1101/816363

**Authors:** Fernando Pires Hartwig, Kate Tilling, George Davey Smith, Deborah A Lawlor, Maria Carolina Borges

**Author notes:** Corresponding author. Postgraduate Program in Epidemiology, Federal University of Pelotas, Pelotas (Brazil) 96020-220.

## Abstract

**Background:** Two-sample Mendelian randomization (MR) allows the use of freely accessible summary association results from genome-wide association studies (GWAS) to estimate causal effects of modifiable exposures on outcomes. Some GWAS adjust for heritable covariables in an attempt to estimate direct effects of genetic variants on the trait of interest. One, both or neither of the exposure GWAS and outcome GWAS may have been adjusted for covariables.

**Methods:** We performed a simulation study comprising different scenarios that could motivate covariable adjustment in a GWAS and analysed real data to assess the influence of using covariable-adjusted summary association results in two-sample MR.

**Results:** In the absence of residual confounding between exposure and covariable, between exposure and outcome, and between covariable and outcome, using covariable-adjusted summary associations for two-sample MR eliminated bias due to horizontal pleiotropy. However, covariable adjustment led to bias in the presence of residual confounding (especially between the covariable and the outcome), even in the absence of horizontal pleiotropy (when the genetic variants would be valid instruments without covariable adjustment). In an analysis using real data from the Genetic Investigation of ANthropometric Traits (GIANT) consortium and UK Biobank, the causal effect estimate of waist circumference on blood pressure changed direction upon adjustment of waist circumference for body mass index.

**Conclusions:** Our findings indicate that using covariable-adjusted summary associations in MR should generally be avoided. When that is not possible, careful consideration of the causal relationships underlying the data (including potentially unmeasured confounders) is required to direct sensitivity analyses and interpret results with appropriate caution.

**Key messages:**

- Summary genetic associations from large genome-wide associations studies (GWAS) have been increasingly used in two-sample Mendelian randomization (MR) analyses.
- Many GWAS adjust for heritable covariates in an attempt to estimate direct genetic effects on the trait of interest.
- In an extensive simulation study, we demonstrate that using covariable-adjusted summary associations may bias MR analyses.
- The bias largely depends on the underlying causal structure, specially the presence of unmeasured common causes between the covariable and the outcome.
- Our findings indicate that using covariable-adjusted summary associations in MR should generally be avoided.

## Introduction

Mendelian randomization (MR) uses genetic variants to assess the influence of modifiable exposures on health outcomes.^1,2^ As germline genetic variants are generally independent of confounding factors and are determined at conception, MR offers a more robust approach to confounding and reverse causation than other methods used in observational studies.^3^

Two-sample MR is an extension to the one-sample MR design, where estimates for the association of genetic variants with exposure and with outcome are derived from different (non-overlapping) samples from the same underlying population.^4^ These estimates are combined to obtain the causal effect estimate of exposure on outcome.^5^ Given that genetic variants typically explain a small proportion of the variation in the exposure of interest, large sample sizes are required for adequately powered MR studies. Therefore, in recent years, two-sample MR has substantially grown in popularity^6^ since it capitalizes on the use of publicly-available summary association results from large genome-wide association studies (GWAS).

In GWAS, estimates for the association of genetic variants with the trait of interest are often conditioned on covariables. As an example, GWAS of waist-to-hip ratio have adjusted estimates for body mass index (BMI),^7^ GWAS of lung function have adjusted estimates for height and stratified analysis by smoking status,^8^ and GWAS of birth weight has excluded pre-term births.^9^ Typically, the aim of conditioning on covariables is estimating the direct effect of genetic variants on the trait (i.e. effects independent of the covariable) or to improve statistical power by reducing residual variance. However, this strategy could introduce bias in GWAS association estimates if the covariable is a collider (or a descendant of a collider) on a pathway linking the genetic variant to the trait of interest. It has previously been shown that conditioning on heritable covariables can bias GWAS association estimates and that the magnitude of this bias is a function of the effect of the genetic variant on the covariable and the correlation structure between the covariable and the trait of interest.^10,11^

The potential issue of adjusting for heritable covariates is illustrated in Figure 1. In this example, a lung function GWAS would be adjusted for smoking in an attempt to identify only the genetic variants with direct effects (i.e., not mediated by smoking) on lung function. In this example, one would hope that the GWAS would identify SNP1 and SNP3 as being associated with lung function (because both have a direct effect on lung function), but not SNP2 (which affects lung function only through its effect on smoking). However, due to the presence of an unmeasured common cause between smoking and lung function (represented by U), smoking is a collider in the path SNP2 → Smoking ← U → Lung function, which links SNP2 and lung function. Because adjustment for a collider opens the path on the collider, SNP2 would be associated with lung function even after adjustment for smoking (eliminating the association would require additional adjustment for U, which is not possible in this example because U is unmeasured). This implies that the adjusted GWAS would be expected to wrongly identify SNP2 as having a direct effect on lung function, and to provide biased estimates for the direct effect of SNP3 on lung function (which would be a combination of the true direct effect and the bias due to collider adjustment). The association between SNP1 and lung function would not be influenced by adjustment for smoking because there is no open path between SNP1 and smoking.

**Figure 1.**
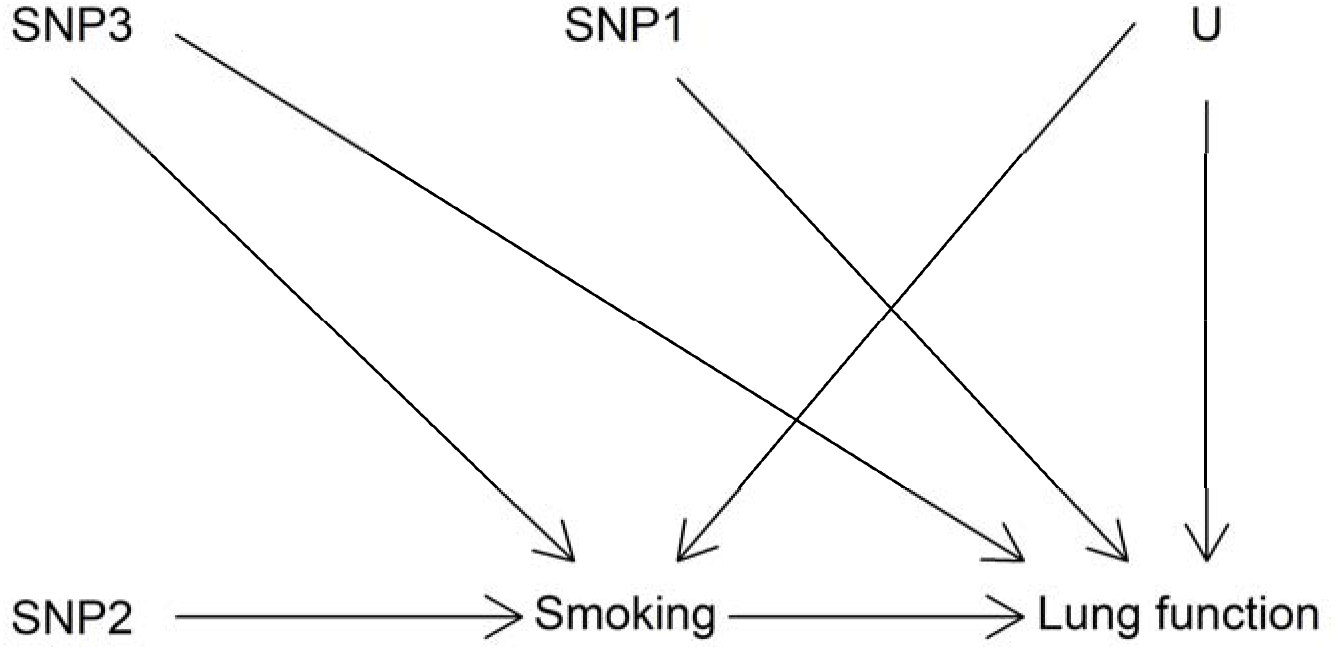
Hypothetical causal structures involving three genetic variants (SNP1-3), smoking, lung function and an unmeasured common cause of the latter two (U).

Several two-sample MR studies have used summary association data from GWAS that have estimated the effect of genetic variants on the trait of interest conditioned on heritable covariables (e.g.^12–16^). The use of such GWAS data might have biased the findings of these MR studies for the reasons outlined above. In addition, it is challenging to predict the impact of such bias since data from two independent GWAS (one for the exposure and other for the outcome) may be used in two-sample MR studies, meaning that the conditional estimates could be restricted to the association of genetic instruments with the exposure, with the outcome, or with both exposure and outcome. Despite there being many published studies using covariable-adjusted summary associations (e.g.^12–16^), few investigations have been made about how this could affect the validity of results (e.g.^12–14,17,18^), particularly in the context of two-sample MR. In this paper we explore how covariable adjustment in GWAS affects two-sample MR findings using simulated and real data in scenarios that could motivate conditioning on a heritable covariable in a GWAS.

## Methods

### Simulation study

We performed a series of simulations to evaluate the consequences of covariable adjustment in MR. We were interested in evaluating situations where genetic variants (*Z*) are marginally associated with both the exposure (*X*) and a covariable (*W*) (Figure 2). This may be because: *Z* causes *W* which causes *X* (Scenarios A, Figure 2); *Z* causes *X* and W independently (Scenarios B and E, Figure 2); *Z* causes *X* which causes *W* (Scenarios C, Figure 2); or *Z* causes an intermediate exposure (*R*) that causes both *X* and *W* (Scenarios D and F, Figure 2). In these situations, the GWAS analyst might decide to adjust for *W* as an attempt to estimate the effect of genetic variants on *X* independent of *W*, thus generating covariable-adjusted summary association results that will be used by two-sample MR analysts to estimate the effect of *X* on an outcome (*Y*), who do not have access to covariable-unadjusted results.

**Figure 2.**
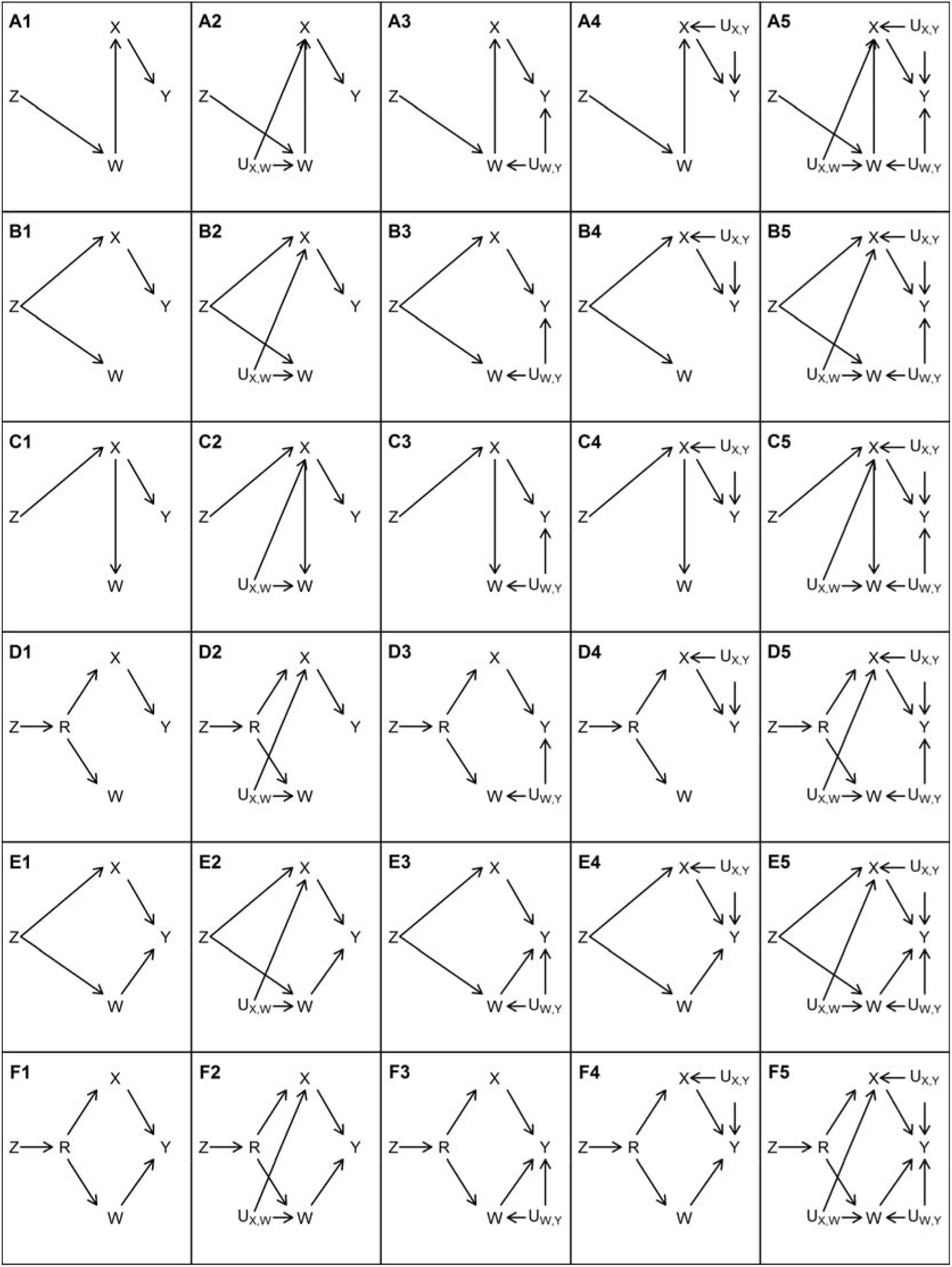
Causal structures that were assessed in the simulation study. ***Z*: genetic instrument; *W*: covariable; *R*: possible direct consequence of *Z*; *X*: exposure; *Y*: outcome: *U*: unmeasured common cause.** Rows represent different causal structures between *Z*, *X*, *W* and *Y* as illustrated by scenarios A (*W* fully mediates the effect of *Z* on *X*), B (*Z* independently affects both *X* and *W*), C (*X* fully mediates the effect of *Z* on *W*), D (effect of *Z* on *X* and *W* is mediated by a common cause *R*), E (same as B except that *W* has a direct effect on *Y*), and F (same as D except that *W* has a direct effect on *Y*). Columns represent different confounding structures between *X*, *W* and *Y* as illustrated by scenarios A1-F1 (no unmeasured confounders other than R), A2-F2 (presence of *X-W* confounder: *U*_*X,W*_), A3-F3 (presence of *W-Y* confounder: *U*_*X,W*_), A4-F4 (presence of *X-Y* confounder *U*_*X,Y*_), and A5-F5 (presence of all three confounders simultaneously).

We performed simulations in two main ways:

- i) All genetic variants have directionally consistent direct effects on the same traits (hereafter referred to as “homogeneous genetic variants”). For example, if there are 10 simulated genetic variants and one of them has positive direct effects on both *X* and *W* (but in no other variable in Figure 2), then all other 9 genetic variants also have positive direct effects on *X* and *W* only. In this scenario, the genetic variants *Z* may have direct effects on *X*, *W* or both, depending on the data-generating model (explained below). These causal structures are shown in Figure 2, where *Z* represents the set of genetic variants.
- ii) Some genetic variants have direct effects only on some traits (hereafter referred to as “heterogeneous genetic variants”). This also varies according to the data-generating model (explained below).

The distinction between situations i) and ii) is further clarified below, after presenting the causal structures underlying the simulations.

Scenarios simulated under situation i) are illustrated in Figure 2. The measured variables are the candidate genetic variants *Z* (which are not necessarily valid instruments for the effect of *X* on *Y*, in which case an MR analysis using *Z* as instruments would be biased), the exposure *X*, the outcome *Y*, the covariable *W*, and (in scenarios D and F) a variable *R* that is a common cause of *X* and *W*. . Although multiple candidate genetic instruments are used in the simulations, Figure 2 only shows one for simplicity (in this scenario, all candidate genetic instruments have the same causal structure, i.e. the homogeneous scenario described above). In all situations, *X* and *W* are genetically correlated (ie, both are marginally associated with *Z*). The aim is to estimate the causal effect of *X* on *Y* using summary GWAS results in a two-sample MR framework. Therefore, there are four possible combinations: no adjustment for *W*; adjusted *Z-X* association but unadjusted *Z-Y* association; unadjusted *Z-X* association but adjusted *Z-Y* association; or both *Z-X* and *Z-Y* associations adjusted for *W*.

Figure 2 depicts the assumed causal structures that we evaluated in the simulations. In scenarios A1-A5, *W* fully mediates the effect of *Z* on *X*. In scenarios B1-B5, *Z* independently affects both *X* and *W* (in other words, *Z* is a confounder of the *X-W* association). In scenarios C1-C5, *X* fully mediates the effect of *Z* on *W*. In scenarios D1-D5, the effect of *Z* on *X* and *W* is mediated by a common cause *R*, so that the effect of *Z* on *W* is correlated with the effect of *Z* on *X*, even though there is no causal effect from *X* to *W* or vice-versa. In all scenarios A1-5 to D1-5, the instrumental variable assumptions hold, so that *Z* is a valid instrument to estimate the causal effect of *X* on *Y*. However, this is not the case in scenarios E1-E5 and F1-F5, which are identical to B1-B5 and D1-D5 (respectively), except that *W* has a direct effect on *Y* (ie, horizontal pleiotropy). In our data generating model (explained in detail in the Supplementary Material), all direct effects are independent on one another. Therefore, the *Z-W* and *Z-X* associations are independent in scenarios E1-E5, meaning that in these scenarios there is horizontal pleiotropy, but the InSIDE (Instrument Strength Independent of Direct Effects) assumption holds. However, in scenarios F1-F5, the InSIDE assumption is violated because Z has an effect on a common cause of *W* and *X*.

In scenarios A1-F1, there are no unmeasured confounders (other than R). To isolate the implications of unmeasured confounders when controlling for *W*, different confounders were included in different scenarios: *X-W* confounder *U_X,W_* in scenarios A2-F2; *W-Y* confounder *U_X,W_* in scenarios A3-F3; *X-Y* confounder *U_X,Y_* in scenarios A4-F4; and finally all three confounders simultaneously in scenarios A5-F5.

To simulate data under situation ii), the same underlying causal structure used for situation i) was used, with the exception that, in all simulations, there were four non-overlapping subgroups of genetic variants: some with direct effects on *X* only, some on *W* only, some on both *X* and *W* (but not *R*), and some on *R* only (in scenarios that include *R*).

It is now easy to clarify the distinction between situations i) and ii). In situation i), all genetic variants have direct effects on *W* only in scenarios A1-A5; on *X* only in scenarios C1-C5; on both *X* and *W* (but in no other variable) in scenarios B1-B5 and E1-E5; on *R* only in scenarios D1-D5 and F1-F5. In situation ii), some genetic variants have direct effects on *X* only, other genetic variants have direct effects on *W* only, and yet other genetic variants have direct effects on both *X* and *W* (but in no other variable) in all scenarios; in scenarios D1-D5 and F1-F5, there is another subset of genetic variants, which have direct effects on *R* only.

All simulations generated data for 40 independent candidate genetic instruments and 100,000 individuals, and the resulting dataset was divided into two halves at random: one was used to estimate instrument-exposure associations, and the other to estimate instrument-outcome associations, thus corresponding to the two-sample MR context. Briefly, for each measured phenotypic variable, all genetic variants with direct effects on the given phenotype were combined into an additive allele score which had an effect set to account for 10% of the variance of the given phenotype. For measured phenotypic and unmeasured variables with direct effects on the given phenotype, direct effects were set to account for 20% of the variance of the given phenotype. A detailed description on how effects were generated is provided in the Supplementary Material. Even though Figure 2 depicts *X* as a having a non-null causal effect on *Y*, all of these scenarios were also simulated for both a null and non-null causal effect from X to Y. A detailed description of the data generating model is provided in the Supplementary Material.

For analyses using unadjusted variant-exposure summary association results (regardless of whether the variant-outcome associations were adjusted for *W*), we selected variants with an unadjusted association with the exposure achieving a *F* statistic of 10 or more to ensure that only variants relatively strongly associated with *X* (which is one of the conditions required for MR to be valid) were included in the analyses. The same logic was applied to analysis using adjusted variant-exposure summary association results.

### Real data example: assessing the causal effect of waist circumference on blood pressure

We conducted an illustrative analysis to explore the impact of covariable adjustment in MR in a real data setting. The exposure of interest was waist circumference (WC), the outcome variables were systolic (SBP) and diastolic blood pressure (DBP), and the covariable was BMI. We selected genetic instruments of unadjusted WC and BMI-adjusted WC from the Genetic Investigation of ANthropometric Traits (GIANT) consortium^7^ and calculated the corresponding instrument-BP summary association results using an interim release of UK Biobank data.^19^ Details on the data sources are provided in the Supplementary Material.

BMI was used as a covariable due to its strong correlation with WC, meaning that variants that affect WC might also affect BMI due to their effect on overall adiposity. Here, we assume that two distinct causal structures (Figure 3) are plausible. In panel A, the genetic variant has a direct effect on a latent variable (which we refer to as adiposity), which manifests itself in measurable constructs such as WC and BMI. Panel B depicts a scenario where the genetic variant has a direct effect on WC rather than on adiposity. Those mechanisms are not mutually exclusive, since different genetic variants can present those different direct effects, or even a single genetic variant can have direct effects on both. Of note, WC, BP, BMI and adiposity are analogous to *X*, *Y*, *W* and *R* (respectively) in Figure 2.

**Figure 3.**
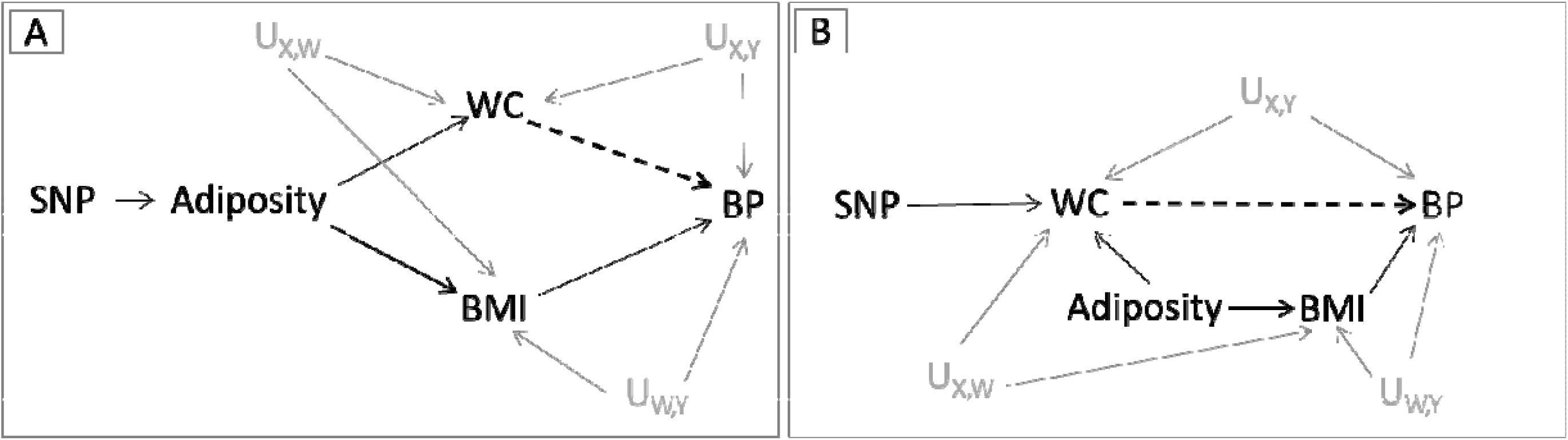
Causal diagrams representing the assumed causal relationships in the two-sample Mendelian randomization analysis of waist circumference (WC) on blood pressure (BP). The genetic instruments (SNPs) are assumed to influence WC by either affecting overall body adiposity (proxied by BMI) [A] or by specifically changing body fat distribution [B]. The grey solid lines represent the effect of confounders between exposure-outcome (U_X,Y_), exposure-covariable (U_X,W_) and covariable-outcome (U_W,Y_). The dashed lines represent the relationship being tested between WC and BP. BMI: body mass index; IV: instrumental variable; SNP: single nucleotide polymorphism; U: unmeasured confounders.

We aimed at replicating scenarios in which the summary association results (either unadjusted or adjusted for the covariable) are already available. Therefore, we selected two sets of WC genetic instruments: one using the unadjusted GWAS results; and another using the BMI-adjusted GWAS. In each case, independent genetic variants were selected as WC instruments if P-value <5×10^−8^. After quality control (described in detail in the Supplementary Materials), 37 single nucleotide polymorphisms (SNPs) and 60 SNPs were retained as genetic instruments for BMI-unadjusted and BMI-adjusted WC, respectively. Prior to analysis, data was harmonised following the steps described in Hartwig et al ^6^ (as detailed in the Supplementary Material). As in the simulation study, four possible combinations were considered: i) unadjusted instrument-WC and unadjusted instrument-BP associations; ii) adjusted instrument-WC and unadjusted instrument-BP associations; iii) unadjusted instrument-WC and adjusted instrument-BP associations; and iv) adjusted instrument-WC and adjusted instrument-BP associations.

### Statistical analyses

Causal effect estimates were obtained using multiplicative random effects inverse-variance weighting (IVW).^5,20^ In the simulation study, coverage and average causal effect estimates were obtained across 5 000 simulated datasets. Coverage was defined the proportion of times that 95% confidence intervals included the true causal effect.

## Results

### Simulation study

Supplementary Table 1 displays the number of select genetic instruments and mean *F* statistic of the instrument-*X* association (among selected instruments), in analyses adjusted and not adjusted for *W*. Since instruments are selected from the exposure GWAS, the set of genetic instruments varies according to the analysis. Taking row 3 of Supplementary Table 1 as an example, on average 35 variants were used for analyses using unadjusted instrument-*X* associations, and 33 variants for analyses using adjusted instrument-*X* association. The mean *F* statistic of 144.7 corresponds to the strength of association between the selected variants (here, 35 on average) and *X* estimated without adjusting for *W*, while the value of 135.3 corresponds to the strength of association between the selected variants (here, 33 on average) and *X* estimated adjusting for *W*. Of note, for homogeneous genetic variants in scenarios A1, A3 and A4, on average the number of selected instruments was 0 upon adjusting the *Z-X* association for *W*. This is expected since conditioning on *W* closes all open paths between *Z* and *X*, without opening new ones (as can be seen in Figure 2). Given that comparing results with and without covariable adjustment is the main goal of this study, we excluded scenarios A1, A3 and A4 from the presentation of results involving homogeneous genetic variants.

Figures 4 and 5 display the bias of the IVW estimate when instrument selection is performed among homogeneous and heterogeneous genetic variants, respectively (coverage of the 95% confidence intervals is shown in Supplementary Figures 1 and 2). Although the absolute bias was generally larger when simulating homogeneous genetic variants, the results were generally in the same direction in both situations.

**Figure 4.**
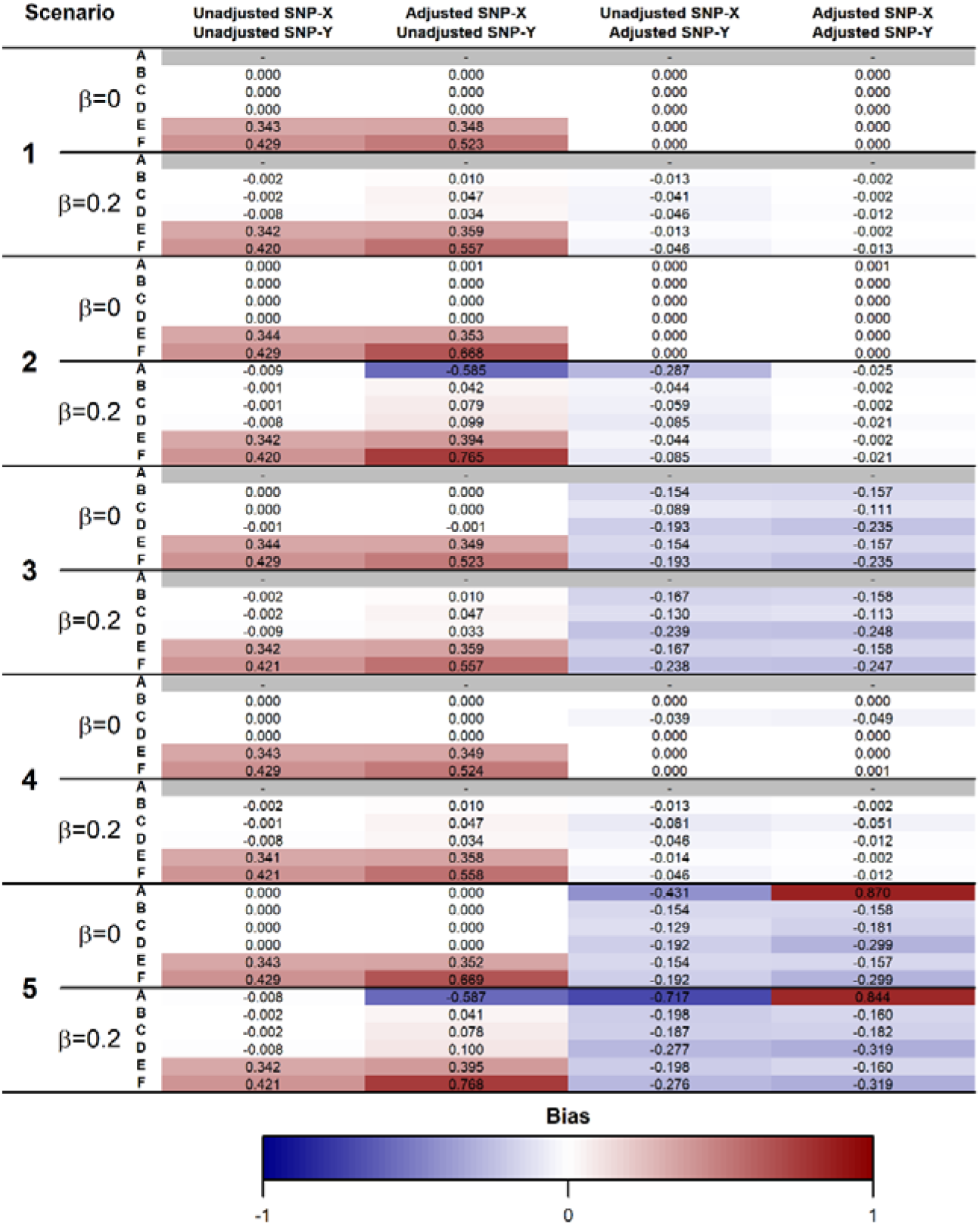
Mean bias across 5,000 simulations of the causal effect estimate using homogeneous genetic instruments. *β*: True causal effect of the exposure (*X*) on the outcome (*Y*). Scenarios A-F assume different causal relationships among *X*, *Y*, the instrument (*Z*), the covariate (*W*) and a common cause of *X* and *W* affected by *Z* (*R*). In scenarios A1-F1, there are no unmeasured confounders. In scenarios A2-F2, there is an unmeasured common cause of *X* and *W*. In scenarios A3-F3, there is an unmeasured common cause of *W* and *Y*. In scenarios A4-F4, there is an unmeasured common cause of *X* and *Y*. In scenarios A5-F5, all these three unmeasured confounders are present. The scenarios are illustrated in Figure 2 and described in detail in the “Simulation study” section. Scenarios where no genetic variants are selected as instruments (because adjustment for *W* results in no open path between *Z* and *X*) are marked in grey.

**Figure 5.**
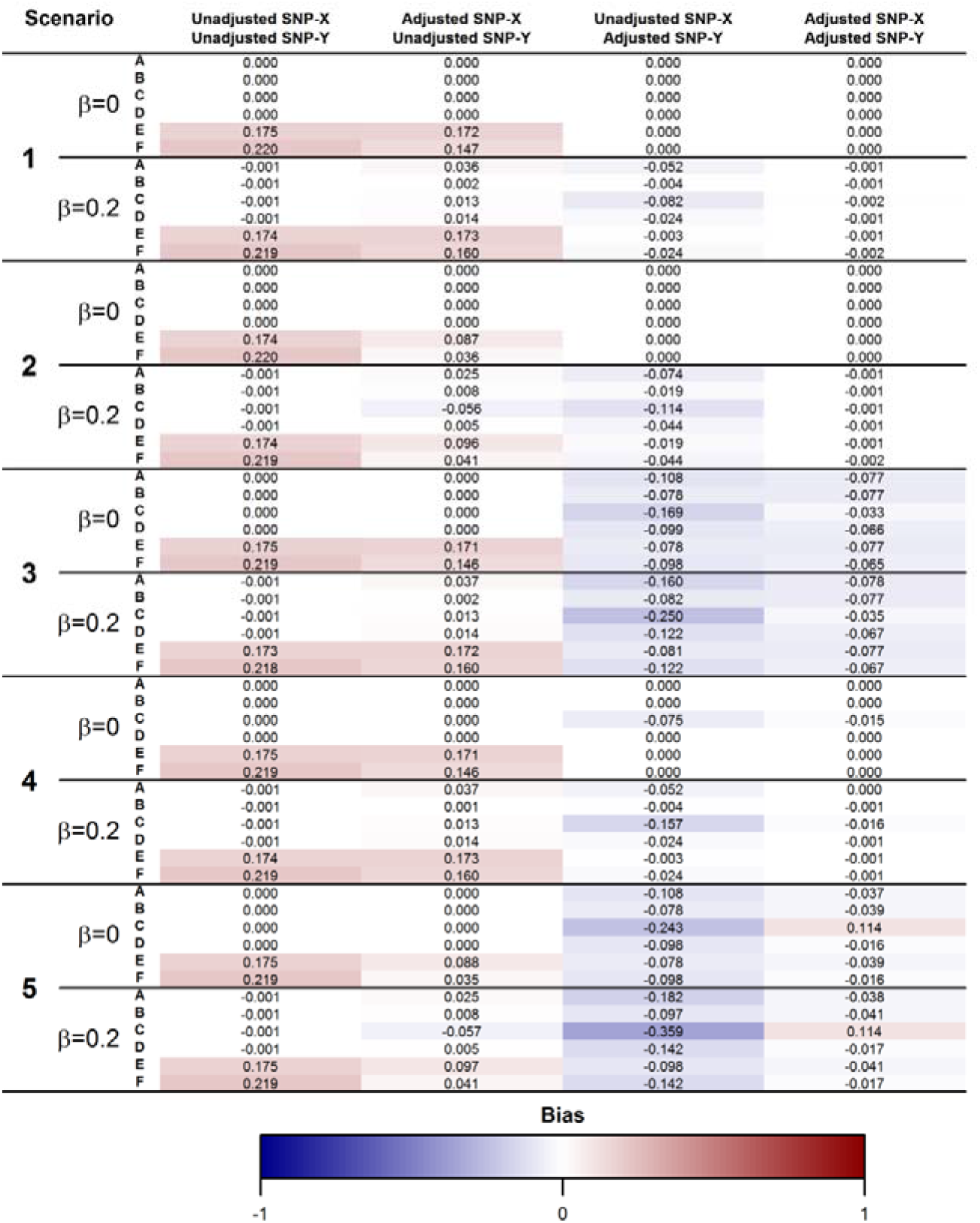
Mean bias across 5,000 simulations of the causal effect estimate using heterogeneous genetic instruments. *β*: True causal effect of the exposure (*X*) on the outcome (*Y*). Scenarios A-F assume different causal relationships among *X*, *Y*, the instrument (*Z*), the covariate (*W*) and a common cause of *X* and *W* affected by *Z* (*R*). In scenarios A1-F1, there are no unmeasured confounders. In scenarios A2-F2, there is an unmeasured common cause of *X* and *W*. In scenarios A3-F3, there is an unmeasured common cause of *W* and *Y*. In scenarios A4-F4, there is an unmeasured common cause of *X* and *Y*. In scenarios A5-F5, all these three unmeasured confounders are present. The scenarios are illustrated in Figure 2 and described in detail in the “Simulation study” section.

In the absence of unmeasured confounders (scenario 1), using adjusted instrument-*Y* associations (especially in combination with adjusted instrument-*X* associations) eliminated bias due to horizontal pleiotropy (scenarios E1 and F1). However, the presence of a *W-Y* confounder (scenario 3) resulted in bias in analyses using covariable-adjusted instrument-*Y* associations even in the absence of horizontal pleiotropy and under the causal null. Moreover, in the presence of a non-null causal effect, the presence of a *X-W* confounder (scenario 2) led to bias in analyses using unadjusted instrument-*X* and adjusted instrument-*Y* associations. Similar trends were observed in the coverage of the 95% confidence intervals.

In cases where *W* does not have a direct effect on *Y* (A-D, and thus none of the variants have horizontal pleiotropic effects), the bias was generally considerably lower in analysis using unadjusted than adjusted instrument-*Y* association estimates. On the other hand, in the presence of horizontal pleiotropy mediated by *W*, analysis using unadjusted instrument-*Y* association estimates presented more bias. This is because all direct effects in our simulations were positive, thus causing the bias introduced by adjusting for *W* (in the presence of unmeasured *W-Y* confounding) to be negative and bias due to horizontal pleiotropy to be positive. In general, bias was larger and coverage lower when the true causal effect was non-zero.

### Real data example

In the unadjusted SNP-exposure and SNP-outcome analysis, each standard unit increase in WC was related to an increase of 0.06 mmHg in SBP (95% CI: −0.01, 0.13) and of 0.12 mmHg in DBP (95% CI: 0.05, 0.19). These effects changed in direction when only adjusting the SNP-outcome association for BMI (SBP: −0.09, 95%CI: −0.17, −0.02; DBP: −0.11, 95%CI: −0.19, −0.04). Effect estimates also changed direction, but were consistent with the null when adjusting only the SNP-exposure association (SBP: −0.06, 95%CI: −0.14, 0.01; DBP: −0.02, 95%CI: −0.09, 0.05) or using both adjusted SNP-exposure and SNP-outcome associations (SBP: −0.04, 95%CI: −0.11, 0.03; DBP: 0.01, 95%CI: −0.06, 0.08) (Figure 6).

**Figure 6.**
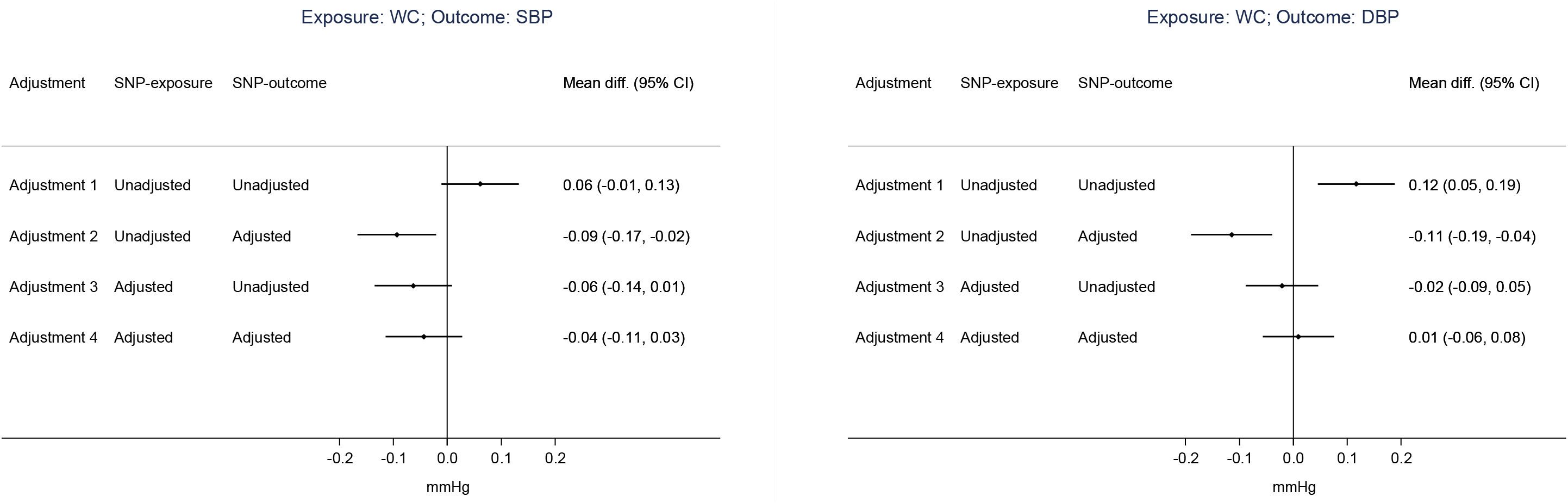
Two-sample Mendelian randomization estimates of the effect of waist circumference (WC) on systolic blood pressure (SBP) or diastolic blood pressure (DBP) for different combinations of adjustments for BMI in SNP-exposure or SNP-outcome association. Effect estimates are expressed as mean difference, and 95% CI, of SBP or DBP (in mmHg) per standard unit increase in WC. 37 SNPs and 60 SNPs were used as instruments for waist circumference unadjusted and adjusted for BMI, respectively.

After excluding SNPs associated with BMI (p < 0.05), results were similar for the adjusted SNP-exposure analyses. The exclusion of BMI-related SNPs dramatically restricted the number of SNPs included in the unadjusted SNP-exposure analyses (two out of 37 SNPs), and, as a result, estimates are highly imprecise (Supplementary Figure 3).

## Discussion

Our results indicate that the impact of covariable adjustment in two-sample MR depends on the causal relations and confounding structure between genetic instruments, exposure, covariable and outcome. In addition, the magnitude and direction of bias will vary depending on whether associations between instrument-exposure, instrument-outcome or both are adjusted for the same covariables. In an analysis using real data from GIANT consortium and UK Biobank, the estimated causal effect of waist circumference on blood pressure changed substantially upon adjustment for BMI.

The strong dependence of the results on the underlying causal structure was expected. In the absence of unobserved common causes (confounders) between exposure-covariable, exposure-outcome and covariable-outcome, covariable adjustment eliminates bias due to horizontal pleiotropy mediated by such covariable (scenarios E and F). However, absence of unobserved confounding is unrealistic in observational studies and is one of the primary motivations for performing MR. In the presence of unobserved confounding, mainly between the covariable and the outcome, covariable adjustment will likely lead to bias even in the absence of horizontal pleiotropy due to collider bias when genetic instruments are marginally associated with the covariable. Therefore, minimising horizontal pleiotropy is generally an invalid justification for using covariable-adjusted summary association results for two-sample MR.

Bias was generally weaker when the true causal effect of the exposure on the outcome is null among several scenarios we evaluated. Indeed, even in the absence of any unmeasured confounding, covariable adjustment may lead to bias when the true causal effect is not null. As an example based on the DAG for scenario A1 in Figure 2, where *W* completely mediates the effect of *Z* on *X* and there is no unmeasured confounding, adjustment for *W* will block the path between *Z* and *X*. Results from two-sample MR using unadjusted *Z-X* but adjusted *Z-Y* estimates will be unbiased if *X* has no causal effect on *Y*, but biased if *X* causes *Y*. Therefore, null results from MR analysis using covariable-adjusted summary association results are generally more reliable (in the sense of being less likely to be a consequence of covariable-adjustment bias) than non-null results, assuming that it is generally unlikely that bias due to using covariable-adjusted summary associations perfectly balances out a given non-null true causal effect. Given that this assumption may be violated (or near violated) depending on the parametrization of the data-generating model, the notion that null results are less likely to be a result of bias should not be interpreted as a general rule.

Using covariable-adjusted instrument-outcome summary associations was also consistently related to more bias compared to using covariable-adjusted instrument-exposure summary associations. The exception was scenarios where there was horizontal pleiotropy. However, since attempting to minimise horizontal pleiotropy via covariable adjustment is generally unjustified due to the likely presence of unmeasured confounders, we are primarily concerned with the scenarios where there is no horizontal pleiotropy.

Results were generally consistent between simulations with homogeneous and heterogeneous instruments. The simulations with homogeneous instruments are useful to isolate the effect of covariable adjustment according to each underlying causal structure linking *Z*, *X*, and *W* illustrated in Figure 2. However, we believe that the most realistic scenarios are the simulations involving heterogeneous instruments; in practice, it is likely that, across the entire genome, there are different subsets of variants affecting *X* via different mechanisms, some of which may or may not influence the covariable *W*.

In an analysis using real data from GIANT consortium and UK Biobank, the estimated causal effect of waist circumference on blood pressure changed substantially upon adjustment for BMI. In this context, interpreting whether covariable adjustment could have affected the validity of results is particularly challenging since adjustment for BMI (as a proxy of overall adiposity) could remove bias due to horizontal pleiotropy and/or introduce bias due to conditioning on a collider (i.e. BMI) in the pathway between instrument-outcome and/or instrument-exposure. As illustrated in Figure 3, the impact of BMI adjustment would differ depending on the mechanism by which genetic variants affect waist circumference since bias could result if genetic variants affect both waist circumference and BMI (Figure 3A), but not if genetic variants affect waist circumference only (Figure 3B). In practice, the net bias from covariable adjustment will depend on the direction and magnitude of confounding and horizontal pleiotropy, the mechanism by which genetic variants affect exposure and whether adjustment was made in instrument-exposure and/or instrument-outcome datasets.

To our knowledge, this is the first study to systematically assess the impact of covariable adjustment in (two-sample) MR. Conditioning on a heritable covariable can introduce bias when the covariable is a collider in the pathway between instrument-exposure and/or instrument-outcome. Collider bias can also be introduced in MR studies in other settings, such as in disease progression studies which include a selected (i.e., case-only) group of individuals^17^.

It is important to emphasise that using covariable-adjusted GWAS summary association results in two-sample MR studies differs from applying multivariable MR (MVMR) on unadjusted GWAS summary association results to estimate the effect of two or more exposures on an outcome.18 The focus of our paper is to investigate bias in two-sample MR due to using covariable-adjusted summary association results, especially in the case when those are the only available option (e.g., when using summary data from GWAS consortia that only performed covariable-adjusted analyses). We are not primarily concerned with whether using covariable-adjusted summary associations reduces bias due to horizontal pleiotropy, even though our results indicate that this is unlikely to be the case in the presence of unmeasured confounding. For this aim, MVMR should be preferred, because MVMR uses genetically predicted variations in both the exposure and covariable(s), and is therefore less susceptible to collider bias.^21^

One of the strengths of our study was that our simulations covered a wide range of scenarios, thus allowing a detailed evaluation of covariable-adjustment bias in a variety of situations. The simulation study was also complemented with a real data example illustrating the strong influence that covariable-adjustment may have not only on the magnitude, but also on the presence and direction of the causal effect estimate. However, any simulation study is a simplification of a reality that is likely to be much more complex. It is impossible to simulate all possible scenarios that might be of relevance to this topic. Moreover, some results, especially quantitative estimates of bias and coverage, are highly dependent on the data-generating model. Therefore, our results should be interpreted qualitatively, as general indications of some of the main aspects related to covariable-adjustment in MR. It was reassuring that some results (as described above) were consistent across scenarios, which indicates that they may apply generally (although not universally) to covariable-adjusted MR.

In conclusion, our findings indicate that using summary association results adjusted for heritable covariables may lead to bias in two-sample MR due to unmeasured confounding. We recommend avoiding adjustment for such covariables in the context of MR. When only covariable-adjusted data is available, it is important to carefully consider the causal structure underlying the research question to understand the potential impact on the results. In such cases, we recommend that researchers refrain from interpreting the causal estimate too literally (which indeed requires parametric assumptions in addition to the core instrumental variable assumptions even in the absence of covariable adjustment). To account for horizontal pleiotropy due to measured covariables, MVMR should be preferred over non-genetic covariable adjustment.

## Funding

KT, GDS, DAL and MCB work in the MRC Integrative Epidemiology Unit at the University of Bristol that receives funding from the UK Medical Research Council [MC_UU_00011/1, MC_UU_00011/3 and MC_UU_00011/6]. DAL’s contribution is also supported by the European Research Council under the European Union’s Seventh Framework Programme [FP/2007-2013] / ERC Grant Agreement [Grant number 669545; DevelopObese] and from the European Union’s Horizon 2020 research and innovation programme under grant agreement No 733206 (LifeCycle). MCB is supported by a Medical Research Council (MRC) Skills Development Fellowship [MR/P014054/1]. DAL is a UK National Institute of Health Research Senior Investigator [NF-SI-0611-10196].

## Acknowledgements

This research has been conducted using the UK Biobank Resource under application number 15825. We would like to thank the participants and researchers from the UK Biobank who contributed or collected data.

## Disclosures

### No competing interests

FPH, KT, GDS, MCB.

### Competing interests

DAL has received funding from several national and international government and charitable organisations, Roche Diagnostics and Medtronic for research unrelated to that presented in this paper.

## Publication Statement

This article has been accepted for publication in the International Journal of Epidemiology, published by Oxford University Press.

