## Supplementary Material for "Bias in two-sample Mendelian randomization when using heritable covariable-adjusted summary associations"

**Simulation study**

**Data generating model**

To assess the consequences of covariable adjustment for two-sample MR, we performed simulations using the following data generating model for $N$ individuals, each indexed by $i$ ($i=1,2,\ldots,N$):

$${Z_{R}}_{i}=\left( \sum_{j=1}^{L} {I_{R}}_{j}{\varphi_{R}}_{j}{G_{i}}_{j} \right)/{\sigma_{ZR}}$$

$${Z_{W}}_{i}=\left( \sum_{j=1}^{L} {I_{W}}_{j}{\varphi_{W}}_{j}{G_{i}}_{j} \right)/{\sigma_{ZW}}$$

$${Z_{X}}_{i}=\left( \sum_{j=1}^{L} {I_{X}}_{j}{\varphi_{X}}_{j}{G_{i}}_{j} \right)/{\sigma_{ZX}}$$

$$R_{i}=\gamma_{R}{Z_{R}}_{i}+{\varepsilon_{R}}_{i}$$

$$W_{i}=\theta_{W,X}{U_{W,X}}_{i}+\theta_{W,Y}{U_{W,Y}}_{i}+\rho_{W}R_{i}+\beta_{W}X_{i}+\gamma_{W}{Z_{W}}_{i}+{\varepsilon_{W}}_{i}$$

$$X_{i}=\theta_{W,X}{U_{W,X}}_{i}+\theta_{X,Y}{U_{X,Y}}_{i}+\rho_{X}R_{i}+\delta_{X}W_{i}+\gamma_{X}{Z_{X}}_{i}+{\varepsilon_{X}}_{i}$$

$Y_{i}=\theta_{W,Y}{U_{W,Y}}_{i}+\theta_{X,Y}{U_{X,Y}}_{i}+\beta_{Y}X_{i}+{\varepsilon_{Y}}_{i}$,

where:

${I_{R}}_{j}$, ${I_{W}}_{j}$ and ${I_{X}}_{j}$ are indicator functions;

${\varphi_{R}}_{j}$, ${\varphi_{W}}_{j}$ and ${\varphi_{X}}_{j}\sim\mathrm{Uniform}(0.1,2)$ , independently;

$G_{ij}\sim\mathrm{Binomial}(2, p_{j})$;

$p_{j}\sim Uniform(0.1,0.9)$;

${U_{W,X}}_{i}$, ${U_{W,Y}}_{i}$, ${U_{X,Y}}_{i}\sim N(0,1)$, independently;

${\varepsilon_{R}}_{i}\sim N(0,\sigma_{\varepsilon_{R}}^{2})$, ${\varepsilon_{W}}_{i}\sim N(0,\sigma_{\varepsilon_{W}}^{2})$, ${\varepsilon_{X}}_{i}\sim N(0,\sigma_{\varepsilon_{X}}^{2})$ and ${\varepsilon_{Y}}_{i}\sim N(0,\sigma_{\varepsilon_{Y}}^{2})$, independently;

$Z_{R}$, $Z_{W}$ and $Z_{X}$ are additive allele scores of $L$ independent genetic instruments on R, W and X, respectively, weighting using the parameters ${\varphi_{R}}_{j}$, ${\varphi_{W}}_{j}$ and ${\varphi_{X}}_{j}$ ($j$=1,2,… $L$). The genetic variables ($G_{j}$) were generated as to mimic bi-allelic SNPs in Hardy-Weinberg equilibrium. Each resulting allele score were divided by its sample standard deviation $(\sigma_{ZR}$, $\sigma_{ZW}$, $\sigma_{ZX}$) to set its variance to one. The values of $\sigma_{\varepsilon_{R}}^{2}$, $\sigma_{\varepsilon_{W}}^{2}$, $\sigma_{\varepsilon_{X}}^{2}$ and $\sigma_{\varepsilon_{Y}}^{2}$ were chosen so that $\mathrm{var}\left( R \right)=\mathrm{var}\left( W \right)=\mathrm{var}\left( X \right)=\mathrm{var}\left( Y \right)$. This all, all variables (except the individual genetic variants) had variance equal to one in all simulations. This allows interpreting the effect of one variable on another as a Pearson correlation coefficient (i.e. standard deviation changes in the dependent variable per standard deviation increment in the independent variable).

$X$, $Y$ and $W$ are the exposure trait, the outcome trait and the covariable, respectively. Here, by covariable we mean a variable that may or may not be adjusted for when estimating instrument-exposure or instrument-outcome associations, or both. $R$ is a potential mediator of the effect of the instruments on $W$ and/or $X$. $U_{W,X}$, $U_{W,Y}$ and $U_{X,Y}$ are unobserved confounders of the $W$-$X$, $W$-$Y$ and $X$-$Y$ associations, respectively, and ${\varepsilon_{R}}_{i}$, ${\varepsilon_{W}}_{i}$, ${\varepsilon_{X}}_{i}$ and ${\varepsilon_{Y}}_{i}$ are random error terms. Figure 1 illustrates the data generating model in a directed acyclic graph (DAG).

Importantly, the parameters $\beta_{W}$ and $\delta_{X}$ were never allowed to be both non-zero in all scenarios. This is to avoid having $X$ and $W$ as both causes and consequences of one another, which would imply that $X$ is a cause of itself (and the same for $W$).

**Simulation contexts**

In all simulations $L$=40 and $N$=100,000, and the resulting dataset was divided into two halves at random: one was used to estimate instrument-exposure associations, and the other to estimate instrument-outcome associations, thus corresponding to the two-sample MR context.

**Situation 1: homogeneous genetic variants**

In all scenarios, ${I_{R}}_{j}={I_{W}}_{j}={I_{X}}_{j}=1$ for all values of $j$. There were six main structures of causal relationships, to which we refer to as A-F. In their most basic forms, these scenarios can be described as follows (parameters not mentioned were set to zero):

- A: $\gamma_{W}=\sqrt{0.1}$, $\delta_{X}$=$\sqrt{0.2}$, $\beta_{Y}\in\{0,0.2\}$.
- B: $\gamma_{W}=\sqrt{0.1}$, $\gamma_{X}=\sqrt{0.1}$, $\beta_{Y}\in\{0,0.2\}$.
- C: $\gamma_{X}=\sqrt{0.1}$, $\beta_{W}$=$\sqrt{0.2}$, $\beta_{Y}\in\{0,0.2\}$.
- D: $\gamma_{R}=\sqrt{0.1}$, $\rho_{W}$=$\sqrt{0.2}$, $\rho_{X}$=$\sqrt{0.2}$, $\beta_{Y}\in\{0,0.2\}$.
- E: $\gamma_{W}=\sqrt{0.1}$, $\gamma_{X}=\sqrt{0.1}$, $\delta_{Y}$=$\sqrt{0.2}$, $\beta_{Y}\in\{0,0.2\}$.
- F: $\gamma_{R}=\sqrt{0.1}$, $\rho_{W}$=$\sqrt{0.2}$, $\rho_{X}$=$\sqrt{0.2}$, $\delta_{Y}$=$\sqrt{0.2}$, $\beta_{Y}\in\{0,0.2\}$.

The above structures of causal relationships were used as a backbone to simulate the following scenarios:

Scenario 1: In this scenario, there were no unobserved confounders, thus corresponding to the causal structures A-F in their most basic form (as described above).

Scenario 2: In this scenario, there was an unobserved common cause of $W$ and $X$. This corresponds to the causal structures A-F, but setting $\theta_{W,X}=\sqrt{0.2}$.

Scenario 3: In this scenario, there was an unobserved common cause of $W$ and $Y$. This corresponds to the causal structures A-F, but setting $\theta_{W,Y}=\sqrt{0.2}$.

Scenario 4: In this scenario, there was an unobserved common cause of $X$ and $Y$. This corresponds to the causal structures A-F, but setting $\theta_{X,Y}=\sqrt{0.2}$.

Scenario 5: This scenario simultaneously included all unobserved common causes present in scenarios 2-4. This corresponds to the causal structures A-F, but setting $\theta_{W,X}=\theta_{W,Y}=\theta_{X,Y}=\sqrt{0.2}$.

**Situation 2: heterogeneous genetic variants**

The scenarios were identical to the ones described above, with the following exceptions. First, $\gamma_{R}=\gamma_{W}=\gamma_{X}=\sqrt{0.1}$ in all scenarios. Second, the indicator functions assumed the value of zero for some values of $j$, in order to generate four subgroups with 10 genetic variants each regarding their direct effects on $R$, $W$ and $X$: i) with direct effects on $X$ only (ie, ${I_{X}}_{j}=1$ and ${I_{R}}_{j}={I_{W}}_{j}=0$); ii) with direct effects on $W$ only (ie, ${I_{W}}_{j}=1$ and ${I_{R}}_{j}={I_{X}}_{j}=0$); iii) with direct effects on both $X$ and $W$ (ie, ${I_{W}}_{j}={I_{X}}_{j}=1$ and ${I_{R}}_{j}=0$); and iv) with direct effects on $R$ only (ie, ${I_{R}}_{j}=1$ and ${I_{W}}_{j}={I_{X}}_{j}=0$). In each simulated dataset, variants were assigned to each subgroup at random.

**Data sources**

**GIANT consortium**

The Genetic Investigation of ANthropometric Traits (GIANT) consortium included up to 210,088 individuals of European ancestry from cohorts genotyped with genome-wide single nucleotide polymorphism (SNP) arrays (n = 57) or Metabochip (n = 44) (1). Genotype data was imputed using CEU population from HapMap as reference. Sample and SNP quality control were undertaken within each study separately (1). Prior to genome-wide association analysis, two sets of adjustments were applied to WC. In the first set, WC was adjusted for age, age^2^, study-specific covariables if necessary, and genomic control inflation factor. In the second set, WC was adjusted for all covariables previously mentioned in addition to BMI. Residuals were calculated for men and women separately and then transformed by the inverse standard normal function. Each directly typed and imputed SNP passing quality control was tested for association with WC (unadjusted or adjusted for BMI) under an additive model in a linear regression framework. Summary data for the association between genetic variants and WC, unadjusted and adjusted for BMI, were downloaded from the GIANT consortium website (<https://www.broadinstitute.org/collaboration/giant/index.php/GIANT_consortium_data_file>).

**UK Biobank**

UK Biobank is a national prospective cohort that recruited more than 500,000 men and women aged 40–69 years between 2006-2010. Until April 2017, genotype data was available for approximately 150,000 participants, which included 50,000 individuals genotyped using UK BiLEVE array and about 100,000 individuals genotyped on the UK Biobank array (2). For imputation, autosome chromosomes were phased using a modified version of the SHAPEIT2 and genotypes were imputed using IMPUTE2 and, as reference panels, UK10K haplotype together with the 1000 Genomes Phase 3 (3). Details on sample and SNP quality control can be found in UK Biobank documentation (2, 3). SBP and DBP were measured using Omron 705 IT electronic blood pressure monitor in two different occasions; these two measures were averaged for each participant. In cases where the largest cuff size was too small for the participant, or if the electronic blood pressure monitor failed to produce a reading, a sphygmomanometer with an inflatable cuff was used in conjunction with a stethoscope (4). For our study, we used individual-level data from approximately 150,000 UK Biobank participants that had genotype data to derive estimates of the association between genetic variants and SBP or DBP, unadjusted and adjusted for BMI, under an additive model in a linear regression framework. All estimates were adjusted for principal components of ancestry.

**Quality control and data harmonisation procedures**

Genetic variants associated with waist circumference (WC) in linkage disequilibrium (R^2^<0.05) were removed using 1000 Genomes reference population and SNP Annotation and Proxy Search (SNAP) tool (5). This resulted in 40 SNPs and 64 SNPs selected as instruments for waist circumference unadjusted and adjusted for body mass index (BMI), respectively. After the exclusion of ambiguous palindromic variants (minor allele frequency ≥30%), 37 SNPs and 60 SNPs were retained as instruments for raw and BMI-adjusted WC, respectively.

Harmonisation of datasets of summary association results of instrument-WC and instrument-blood pressure associations was performed following the steps described in Hartwig et al (6). This included: (A) ensuring that all variants in dataset 1 (GIANT) were associated with WC in the same direction (in this case, all effect alleles relate to exposure increasing alleles); (B) ensuring that datasets 1 (GIANT) and 2 (UK Biobank) were identically coded regarding the exposure increasing allele; (C) recoding palindromic SNPs based on the effect allele frequency and removing the ambiguous ones; (D) checking the correlation of effect allele frequencies between datasets 1 (GIANT) and 2 (UK Biobank) before (~0.11) and after harmonization (~0.98).

**Sensitivity analysis**

As a negative control for the impact of bias due to adjustment for BMI in the association between WC and SBP or DBP, we conducted a sensitivity analysis excluding SNPs associated with BMI at p < 0.05, which resulted in only two SNPs being included as instruments for WC unadjusted for BMI (in contrast to 37 SNPs in the main analysis) and in 45 SNPs being included as instruments for WC adjusted for BMI (in contrast to 60 SNPs in the main analysis). Due to the small number of instruments eligible for the analyses of WC unadjusted for BMI (when excluding BMI-associated SNPs), Mendelian randomization estimates were calculated by combining SNP-specific Wald ratios (SNP-outcome beta/SNP-exposure beta) in a fixed-effect meta-analysis framework (using inverse variance weights) (7).

**Supplementary Tables**

**Supplementary Table 1. Mean number of selected genetic instrumental variables (IVs) and mean F-statistic of the IV-**$\boldsymbol{X}$ **association for each simulation scenario.**

| **Scenario** | **IVs** | **Causal** | $\boldsymbol{\beta}_{\boldsymbol{Y}}$ | **Number of selected IVs** | | **Mean** $\boldsymbol{F}$ **statistic** | |
| --- | --- | --- | --- | --- | --- | --- | --- |
|  |  | **structure** |  | **Unadjusted** | **Adjusted for** $\boldsymbol{W}$ | **Unadjusted** | **Adjusted for** $\boldsymbol{W}$ |
| 1 | Homogeneous | A | 0 | 26 | 0 | 38.3 | - |
| 1 | Homogeneous | A | 0.2 | 26 | 0 | 38.3 | - |
| 1 | Homogeneous | B | 0 | 35 | 33 | 144.7 | 135.3 |
| 1 | Homogeneous | B | 0.2 | 35 | 33 | 144.7 | 135.2 |
| 1 | Homogeneous | C | 0 | 35 | 34 | 144.6 | 118.7 |
| 1 | Homogeneous | C | 0.2 | 35 | 34 | 144.9 | 118.9 |
| 1 | Homogeneous | D | 0 | 26 | 22 | 38.2 | 28.9 |
| 1 | Homogeneous | D | 0.2 | 26 | 22 | 38.4 | 29.1 |
| 1 | Homogeneous | E | 0 | 35 | 33 | 144.7 | 135.3 |
| 1 | Homogeneous | E | 0.2 | 35 | 33 | 144.6 | 135.2 |
| 1 | Homogeneous | F | 0 | 26 | 22 | 38.3 | 29.0 |
| 1 | Homogeneous | F | 0.2 | 26 | 22 | 38.2 | 28.9 |
| 1 | Heterogeneous | A | 0 | 27 | 19 | 293.3 | 344.8 |
| 1 | Heterogeneous | A | 0.2 | 27 | 19 | 294.1 | 344.6 |
| 1 | Heterogeneous | B | 0 | 19 | 18 | 272.8 | 267.6 |
| 1 | Heterogeneous | B | 0.2 | 19 | 18 | 273.6 | 268.4 |
| 1 | Heterogeneous | C | 0 | 19 | 25 | 272.8 | 147.1 |
| 1 | Heterogeneous | C | 0.2 | 19 | 25 | 273.1 | 147.0 |
| 1 | Heterogeneous | D | 0 | 27 | 31 | 224.1 | 171.5 |
| 1 | Heterogeneous | D | 0.2 | 27 | 31 | 223.9 | 171.4 |
| 1 | Heterogeneous | E | 0 | 19 | 18 | 273.0 | 267.6 |
| 1 | Heterogeneous | E | 0.2 | 19 | 18 | 273.3 | 268.0 |
| 1 | Heterogeneous | F | 0 | 27 | 31 | 223.9 | 171.0 |
| 1 | Heterogeneous | F | 0.2 | 27 | 31 | 223.8 | 171.1 |
| 2 | Homogeneous | A | 0 | 26 | 15 | 38.2 | 20.0 |
| 2 | Homogeneous | A | 0.2 | 26 | 15 | 38.3 | 20.1 |
| 2 | Homogeneous | B | 0 | 35 | 30 | 144.7 | 118.2 |
| 2 | Homogeneous | B | 0.2 | 35 | 30 | 144.7 | 118.2 |
| 2 | Homogeneous | C | 0 | 35 | 34 | 144.8 | 127.9 |
| 2 | Homogeneous | C | 0.2 | 35 | 34 | 144.7 | 127.5 |
| 2 | Homogeneous | D | 0 | 26 | 18 | 38.3 | 22.3 |
| 2 | Homogeneous | D | 0.2 | 26 | 18 | 38.3 | 22.4 |
| 2 | Homogeneous | E | 0 | 35 | 30 | 144.9 | 118.8 |
| 2 | Homogeneous | E | 0.2 | 35 | 30 | 144.5 | 118.2 |
| 2 | Homogeneous | F | 0 | 26 | 18 | 38.2 | 22.3 |
| 2 | Homogeneous | F | 0.2 | 26 | 18 | 38.2 | 22.3 |
| 2 | Heterogeneous | A | 0 | 27 | 25 | 293.7 | 337.5 |
| 2 | Heterogeneous | A | 0.2 | 27 | 25 | 293.8 | 337.2 |
| 2 | Heterogeneous | B | 0 | 19 | 23 | 273.4 | 204.7 |
| 2 | Heterogeneous | B | 0.2 | 19 | 23 | 273.6 | 204.9 |
| 2 | Heterogeneous | C | 0 | 19 | 26 | 273.3 | 212.9 |
| 2 | Heterogeneous | C | 0.2 | 19 | 26 | 273.6 | 213.2 |
| 2 | Heterogeneous | D | 0 | 27 | 33 | 223.7 | 174.7 |
| 2 | Heterogeneous | D | 0.2 | 27 | 33 | 223.9 | 174.2 |
| 2 | Heterogeneous | E | 0 | 19 | 23 | 273.3 | 204.7 |
| 2 | Heterogeneous | E | 0.2 | 19 | 23 | 272.9 | 205.1 |
| 2 | Heterogeneous | F | 0 | 27 | 33 | 224.2 | 174.2 |
| 2 | Heterogeneous | F | 0.2 | 27 | 33 | 223.7 | 174.1 |
| 3 | Homogeneous | A | 0 | 26 | 0 | 38.3 | - |
| 3 | Homogeneous | A | 0.2 | 26 | 0 | 38.2 | - |
| 3 | Homogeneous | B | 0 | 35 | 33 | 144.8 | 135.3 |
| 3 | Homogeneous | B | 0.2 | 35 | 33 | 144.6 | 135.1 |
| 3 | Homogeneous | C | 0 | 35 | 34 | 144.8 | 118.8 |
| 3 | Homogeneous | C | 0.2 | 35 | 34 | 144.5 | 118.7 |
| 3 | Homogeneous | D | 0 | 26 | 22 | 38.2 | 28.9 |
| 3 | Homogeneous | D | 0.2 | 26 | 22 | 38.2 | 28.9 |
| 3 | Homogeneous | E | 0 | 35 | 33 | 144.5 | 134.9 |
| 3 | Homogeneous | E | 0.2 | 35 | 33 | 144.8 | 135.4 |
| 3 | Homogeneous | F | 0 | 26 | 22 | 38.2 | 28.9 |
| 3 | Homogeneous | F | 0.2 | 26 | 22 | 38.3 | 29.0 |
| 3 | Heterogeneous | A | 0 | 27 | 19 | 293.3 | 345.2 |
| 3 | Heterogeneous | A | 0.2 | 27 | 19 | 293.1 | 344.9 |
| 3 | Heterogeneous | B | 0 | 19 | 18 | 273.1 | 267.9 |
| 3 | Heterogeneous | B | 0.2 | 19 | 18 | 272.9 | 267.5 |
| 3 | Heterogeneous | C | 0 | 19 | 25 | 273.8 | 147.1 |
| 3 | Heterogeneous | C | 0.2 | 19 | 25 | 273.6 | 147.0 |
| 3 | Heterogeneous | D | 0 | 27 | 31 | 223.9 | 171.2 |
| 3 | Heterogeneous | D | 0.2 | 27 | 31 | 223.7 | 170.8 |
| 3 | Heterogeneous | E | 0 | 19 | 18 | 273.2 | 267.9 |
| 3 | Heterogeneous | E | 0.2 | 19 | 18 | 273.2 | 267.9 |
| 3 | Heterogeneous | F | 0 | 27 | 31 | 223.8 | 171.0 |
| 3 | Heterogeneous | F | 0.2 | 27 | 31 | 223.9 | 171.4 |
| 4 | Homogeneous | A | 0 | 26 | 0 | 38.1 | - |
| 4 | Homogeneous | A | 0.2 | 26 | 0 | 38.3 | - |
| 4 | Homogeneous | B | 0 | 35 | 33 | 144.5 | 135.2 |
| 4 | Homogeneous | B | 0.2 | 35 | 33 | 144.8 | 135.5 |
| 4 | Homogeneous | C | 0 | 35 | 34 | 144.6 | 118.6 |
| 4 | Homogeneous | C | 0.2 | 35 | 34 | 144.7 | 118.7 |
| 4 | Homogeneous | D | 0 | 26 | 22 | 38.3 | 29.0 |
| 4 | Homogeneous | D | 0.2 | 26 | 22 | 38.2 | 28.9 |
| 4 | Homogeneous | E | 0 | 35 | 33 | 144.9 | 135.5 |
| 4 | Homogeneous | E | 0.2 | 35 | 33 | 144.6 | 135.1 |
| 4 | Homogeneous | F | 0 | 26 | 22 | 38.3 | 29.0 |
| 4 | Homogeneous | F | 0.2 | 26 | 22 | 38.2 | 28.9 |
| 4 | Heterogeneous | A | 0 | 27 | 19 | 292.8 | 344.9 |
| 4 | Heterogeneous | A | 0.2 | 27 | 19 | 293.9 | 345.5 |
| 4 | Heterogeneous | B | 0 | 19 | 18 | 273.4 | 268.1 |
| 4 | Heterogeneous | B | 0.2 | 19 | 18 | 272.7 | 267.4 |
| 4 | Heterogeneous | C | 0 | 19 | 25 | 273.1 | 147.2 |
| 4 | Heterogeneous | C | 0.2 | 19 | 25 | 273.1 | 147.3 |
| 4 | Heterogeneous | D | 0 | 27 | 31 | 223.8 | 171.1 |
| 4 | Heterogeneous | D | 0.2 | 27 | 31 | 223.9 | 171.1 |
| 4 | Heterogeneous | E | 0 | 19 | 18 | 273.1 | 267.7 |
| 4 | Heterogeneous | E | 0.2 | 19 | 18 | 273.1 | 267.8 |
| 4 | Heterogeneous | F | 0 | 27 | 31 | 223.8 | 171.1 |
| 4 | Heterogeneous | F | 0.2 | 27 | 31 | 224.2 | 171.2 |
| 5 | Homogeneous | A | 0 | 26 | 15 | 38.2 | 20.1 |
| 5 | Homogeneous | A | 0.2 | 26 | 15 | 38.3 | 20.1 |
| 5 | Homogeneous | B | 0 | 35 | 30 | 144.9 | 118.7 |
| 5 | Homogeneous | B | 0.2 | 35 | 30 | 144.6 | 118.4 |
| 5 | Homogeneous | C | 0 | 35 | 34 | 144.8 | 127.9 |
| 5 | Homogeneous | C | 0.2 | 35 | 34 | 144.7 | 127.7 |
| 5 | Homogeneous | D | 0 | 26 | 18 | 38.2 | 22.4 |
| 5 | Homogeneous | D | 0.2 | 26 | 18 | 38.3 | 22.3 |
| 5 | Homogeneous | E | 0 | 35 | 30 | 144.5 | 118.0 |
| 5 | Homogeneous | E | 0.2 | 35 | 30 | 144.8 | 118.2 |
| 5 | Homogeneous | F | 0 | 26 | 18 | 38.3 | 22.3 |
| 5 | Homogeneous | F | 0.2 | 26 | 18 | 38.4 | 22.4 |
| 5 | Heterogeneous | A | 0 | 27 | 25 | 293.5 | 338.1 |
| 5 | Heterogeneous | A | 0.2 | 27 | 25 | 293.2 | 337.5 |
| 5 | Heterogeneous | B | 0 | 19 | 23 | 273.0 | 204.6 |
| 5 | Heterogeneous | B | 0.2 | 19 | 23 | 273.4 | 205.0 |
| 5 | Heterogeneous | C | 0 | 19 | 26 | 273.2 | 213.6 |
| 5 | Heterogeneous | C | 0.2 | 19 | 26 | 272.9 | 213.3 |
| 5 | Heterogeneous | D | 0 | 27 | 33 | 224.1 | 174.6 |
| 5 | Heterogeneous | D | 0.2 | 27 | 33 | 223.9 | 174.5 |
| 5 | Heterogeneous | E | 0 | 19 | 23 | 273.2 | 204.5 |
| 5 | Heterogeneous | E | 0.2 | 19 | 23 | 273.2 | 204.6 |
| 5 | Heterogeneous | F | 0 | 27 | 33 | 224.0 | 174.8 |
| 5 | Heterogeneous | F | 0.2 | 27 | 33 | 223.7 | 174.3 |

**Supplementary Figures**

**Supplementary Figure 1. Mean coverage of 95% confidence intervals across 5,000 simulations using heterogeneous genetic instruments.**

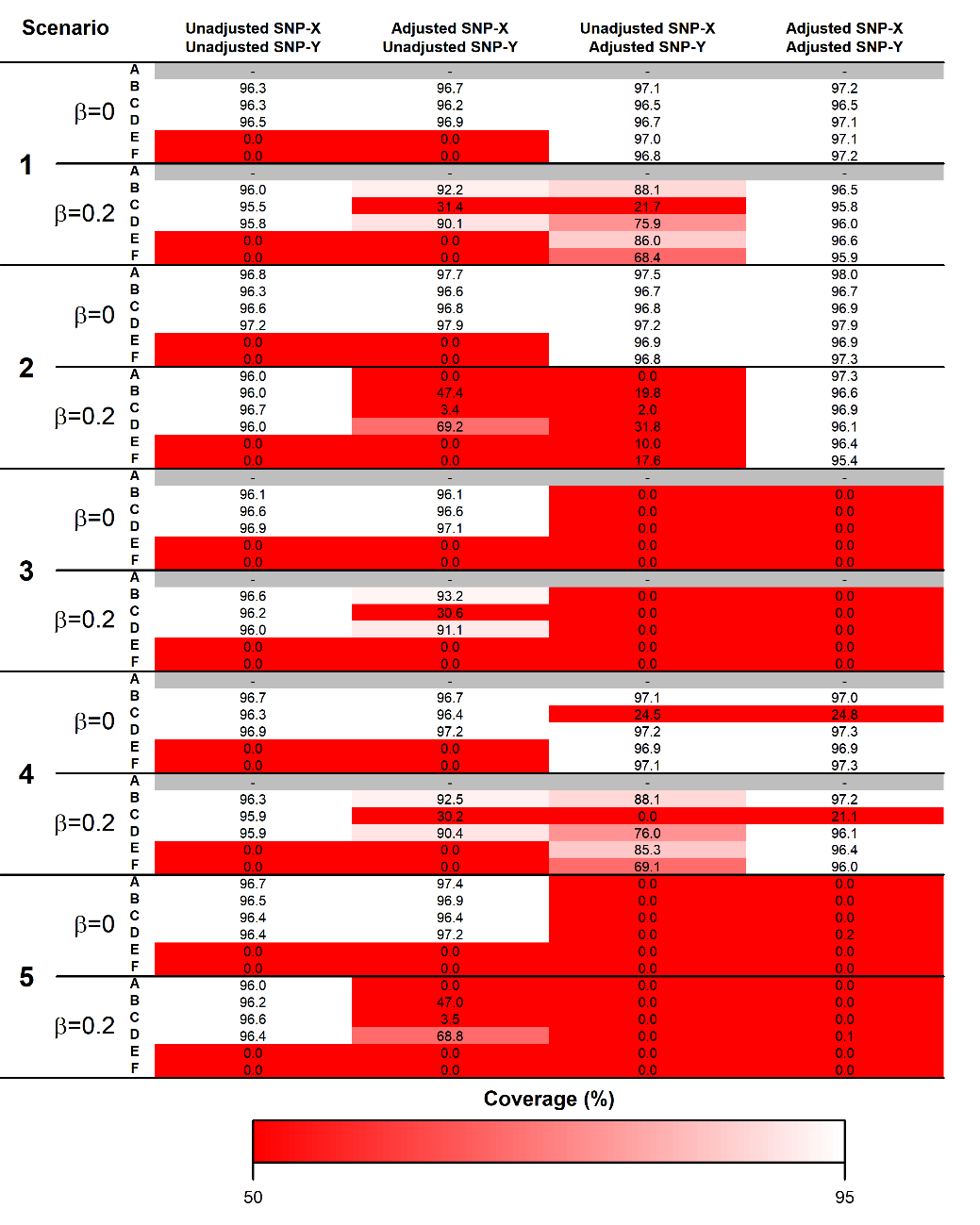

$\beta:$ True causal effect of the exposure ($X$) on the outcome ($Y$).

Scenarios A-F assume different causal relationships among $X$, $Y$, the instrument ($Z$), the covariate ($W$) and a common cause of $X$ and $W$ affected by $Z$ ($R$). In scenarios A1-F1, there are no unmeasured confounders. In scenarios A2-F2, there is an unmeasured common cause of $X$ and $W$. In scenarios A3-F3, there is an unmeasured common cause of $W$ and $Y$. In scenarios A4-F4, there is an unmeasured common cause of $X$ and $Y$. In scenarios A5-F5, all these three unmeasured confounders are present. The scenarios are illustrated in Figure 1 and described in detail in the “Simulation study” section. Scenarios where no genetic variants are selected as instruments (because adjustment for $W$ results in no open path between $Z$ and $X$) are marked in grey.

**Supplementary Figure 2. Mean coverage of 95% confidence intervals across 5,000 simulations using heterogeneous genetic instruments.**

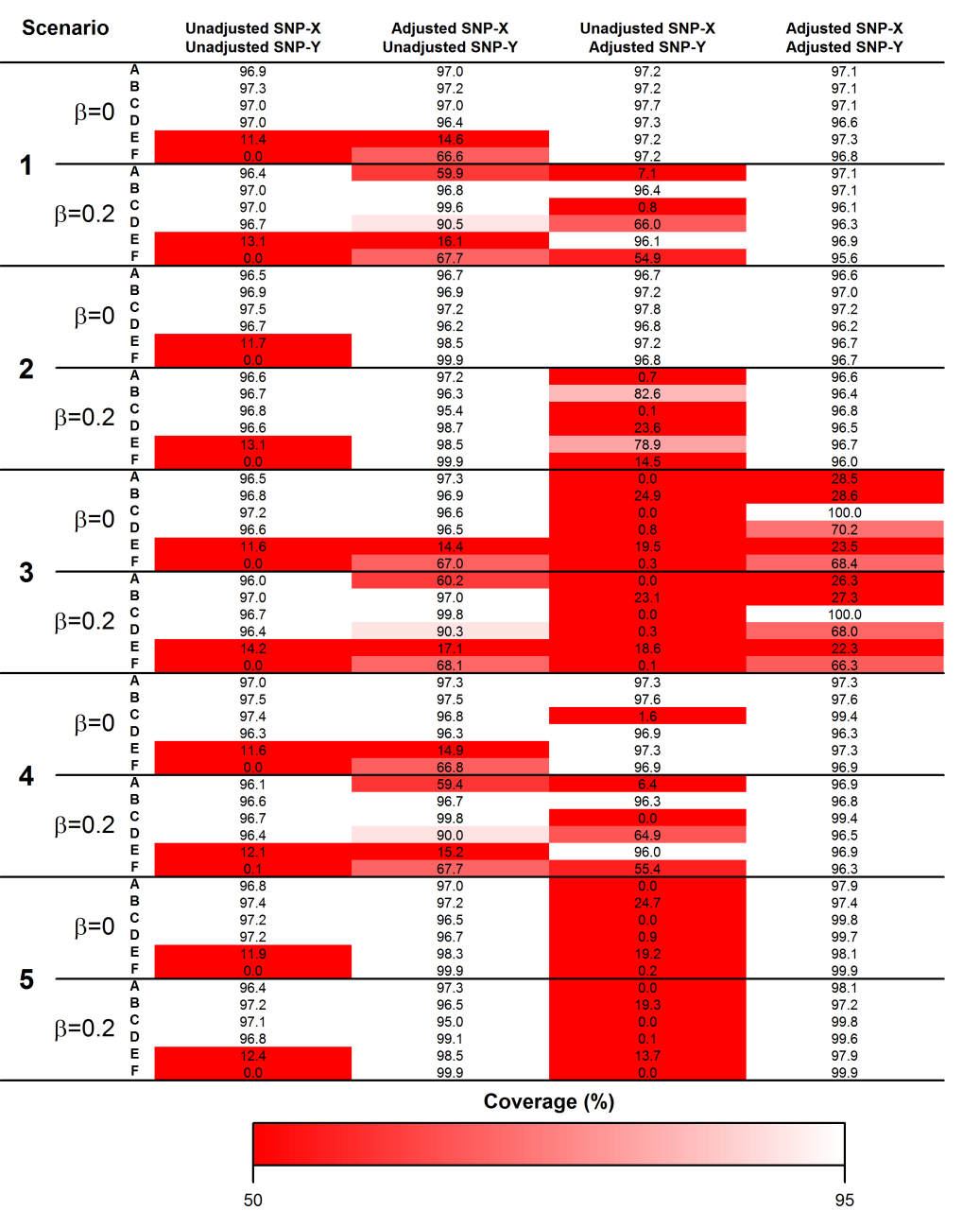

𝛽: True causal effect of the exposure (𝑋) on the outcome (𝑌).

Scenarios A-F assume different causal relationships among 𝑋, 𝑌, the instrument (𝑍), the covariate (𝑊) and a common cause of 𝑋 and 𝑊 affected by 𝑍 (𝑅). In scenarios A1-F1, there are no unmeasured confounders. In scenarios A2-F2, there is an unmeasured common cause of 𝑋 and 𝑊. In scenarios A3-F3, there is an unmeasured common cause of 𝑊 and 𝑌. In scenarios A4-F4, there is an unmeasured common cause of 𝑋 and 𝑌. In scenarios A5-F5, all these three unmeasured confounders are present. The scenarios are illustrated in Figure 1 and described in detail in the “Simulation study” section.

**Supplementary Figure 3. Two-sample Mendelian randomization estimates of the effect of waist circumference (WC) on systolic blood pressure (SBP) or diastolic blood pressure (DBP) including (BMI SNPs: yes) or removing (BMI SNPs: no) SNPs associated with BMI.**

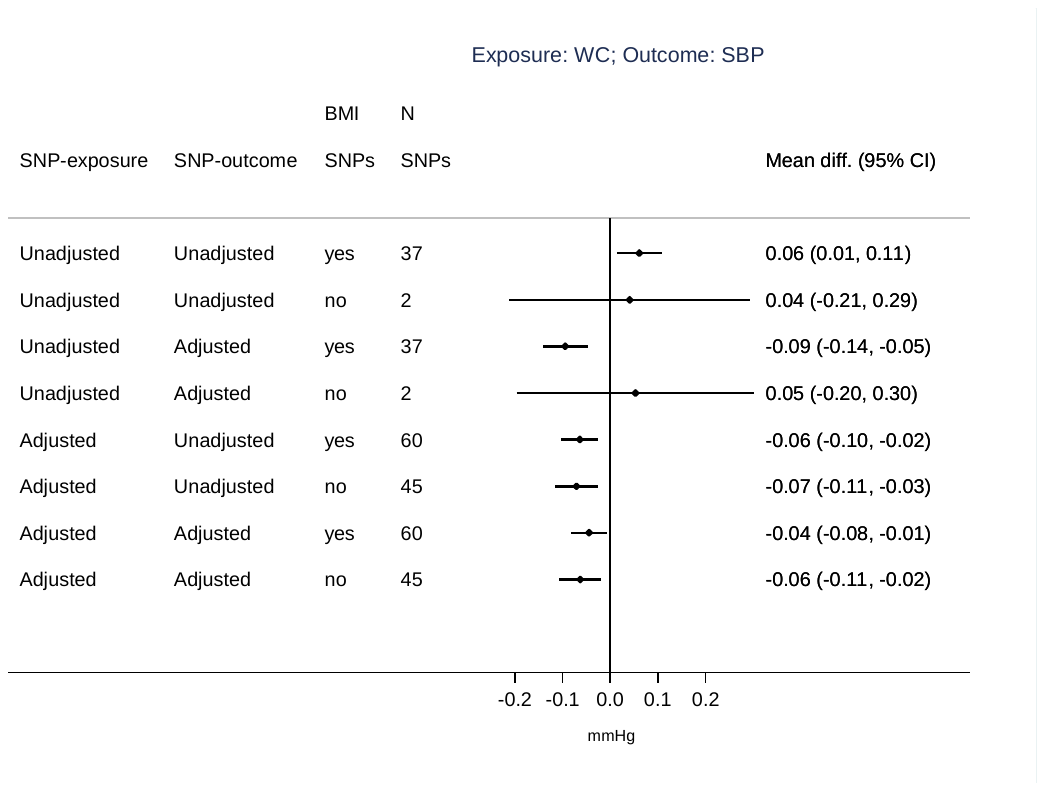

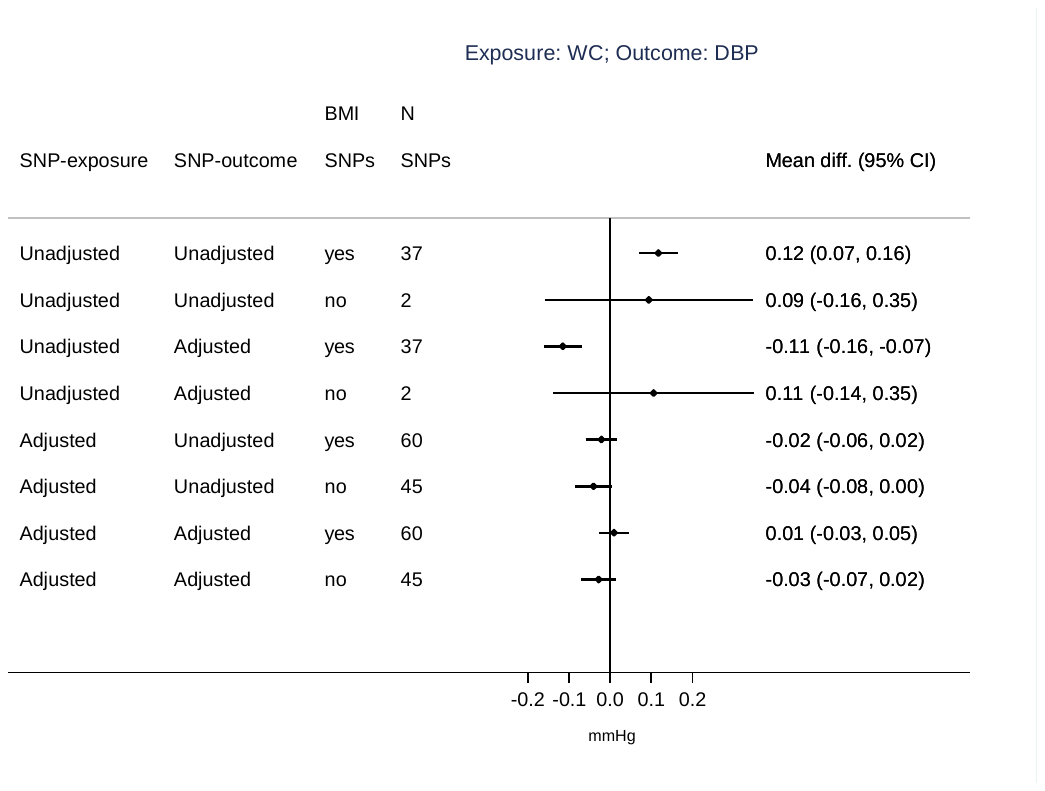

Effect estimates are expressed as mean difference, and 95% CI, of SBP or DBP (in mmHg) per standard unit increase in WC. 37 SNPs and 60 SNPs were used as instruments for WC unadjusted and adjusted for BMI, respectively. After removing instruments associated with BMI (p-value < 0.05), 2 SNPs and 45 SNPs were retained as instruments for unadjusted and BMI-adjusted WC, respectively.
